# Cross-omics profiling reveals cortical neuronal dysfunction following neonatal intraventricular haemorrhage that is attenuated by decorin treatment

**DOI:** 10.64898/2026.09.07.749813

**Authors:** Benjamin J. Hewitt, Maria Garcia Bonilla, James P. McAllister, Simran Pahuja, Grant A. Pellowe, Mackenzie Newman, Lauren Roberts, James A. Roberts, Philip Kitchen, Roslyn M. Bill, Lisa J. Hill, Hannah F. Botfield

## Abstract

Neonatal intraventricular haemorrhage (IVH) is a form of haemorrhagic stroke which is most common in premature infants and can cause mortality and lifelong disability. The reaction of brain tissue to extraneous blood is known to trigger several complications including post-haemorrhagic hydrocephalus (PHH), furthering the likelihood of neurological disability and death. The molecular response of the neuron-rich cortex to blood within the first 24 hours of injury, and how this may precipitate secondary brain injury or neurodegeneration, remains poorly defined. This lack of understanding may reduce the development of novel therapeutics aiming to limit the injury caused by neonatal IVH. Here, we utilised a proteome and transcriptome cross-omics approach to characterise cortical responses to blood in a neonatal mouse model of IVH to improve understanding of the effects and drivers of IVH-induced brain injury. The pleiotropic proteoglycan decorin, a well-characterised modulator of the inflammatory cytokine transforming growth factor beta 1, was also studied to identify if it can have any beneficial effects in the cortex. Cross-omics analysis between the transcriptome and proteome highlighted ferroptosis-associated signatures as a key post-IVH cortical injury mechanism. IVH caused a marked depletion of cortical neurons, neurofilaments, and synaptic vesicle proteins, observed through transcriptomics, immunostaining, and proteomics, which were attenuated by decorin treatment. Decorin also produced an enhanced phagocytic response potentially through activation of microglia. These findings highlight the vulnerability of cortical neurons to early injury following IVH in neonates and support decorin as a potential neuroprotective agent. They also suggest that ferroptosis-associated pathways may contribute to early cortical injury and represent a potential therapeutic target in this context.

## 2. Introduction

Intraventricular haemorrhage (IVH) is a life-threatening form of haemorrhagic stroke characterised by bleeding into the cerebrospinal fluid (CSF)-filled ventricles of the brain. IVH mainly affects very premature neonates (< 32 weeks gestation) with birthweight under 1000 g (Kadri et al., 2006; Piscopo et al., 2025), born prior to the natural regression of the germinal matrix, a highly vascular and structurally vulnerable region of the brain. In these infants, vascular fragility can precipitate germinal matrix haemorrhage (GMH), with extension into the lateral ventricles termed IVH (Towbin, 1968). Approximately 34% of premature infants experience some grade of IVH (Lai et al., 2022), with up to 41% of these suffering from complications as a result (Kadri et al., 2006). In neonates, IVH severity has been shown to strongly correlate with neurodevelopmental impairment, including motor and cognitive defects (Mukerji et al., 2015), and with increased childhood hospitalisation due to other comorbidities (Kaur et al., 2020).

In both infant and adult IVH, the mechanical mass effect of blood on the parenchyma is compounded by tissue damage due to the inflammatory nature of blood products on the parenchyma, ependyma, and dural tissues (Madangarli et al., 2019). Complications such as post-haemorrhagic hydrocephalus (PHH) are common, wherein intracranial pressure (ICP) is markedly increased by a reduction in CSF outflow and increase in CSF secretion (Lolansen et al., 2022). Conventional treatment regimens for neonatal IVH are surgical in nature, attempting to reduce ICP and treating resultant symptoms (including seizures, bradycardia, and respiratory distress). Up to 60% of survivors of severe IVH develop PHH and 25% require treatment by surgical intervention (e.g shunting) to lower ICP (Vassilyadi et al., 2009). However, existing treatment approaches do not tackle the effects that blood-laden CSF has on the brain tissue.

The underlying pathogenic mechanisms which follow IVH and how these may influence secondary brain injury and neurodevelopmental impairment in neonates is not fully understood, though iron from free blood is known to play a role in the development of PHH (Strahle et al., 2014) and lysed blood has been shown to directly cause extracellular matrix (ECM) dysfunction in the absence of mass effect (Hewitt et al., 2026). Neurological function is critically dependent upon the cortex, being the site of specialised layers of neurons with both afferent and efferent functions. In addition, the generation of neurons from progenitor cells located in the periventricular region (the ventricular/subventricular zone) and their localisation is a crucial part of neurodevelopment, and thus any injury to the cortex either *in utero* or in the early postnatal days may be expected to have substantial impact on neurological function (Campbell, 2005). This is highlighted by the high incidence of neurological disability after neonatal IVH and the marked reduction in quality of life for survivors of IVH (Rees et al., 2025).

Despite this vulnerability, the cortical response to IVH remains relatively unexplored compared with the ependyma and periventricular white matter, despite exposure to the same cytotoxic and pro-inflammatory blood products within the CSF after circulation through the ventricular system. Additionally, the use of a suitably targeted and delivered pharmacological therapy within an early window after IVH may reduce cortical tissue damage, prevent neuronal death, and thus improve neurological outcomes and reduce complications such as cerebral palsy. The development of a non-surgical treatment to improve outcomes after neonatal IVH is a key research gap with the potential to reduce mortality and lifelong morbidity which can often follow from this devastating condition.

Here, we describe a novel cross-omics approach to improving understanding of the early response of cortical brain tissue to neonatal IVH, utilising a neonatal IVH model with downstream RNA-sequencing, proteomics, and immunofluorescence staining to identify key changes in mouse brains 24 hours after IVH at post-natal day 4. We have previously shown that increased transforming growth factor beta 1 (TGF-β1) expression after haemorrhagic stroke is associated with ECM dysfunction and PHH development (Hewitt et al., 2025), and thus we also trialled the TGF-β1 antagonist decorin to examine its possible effects on the cortex in neonatal IVH. Decorin is an endogenous proteoglycan which has previously been trialled in juvenile communicating hydrocephalus to excellent effect in pre-clinical trials; lowering the incidence of ventriculomegaly and reducing ECM deposition whilst supressing the inflammatory response via reductions in TGF-β1 expression and pSMAD2/3 signalling (Botfield et al., 2013).

Both over-representation analysis and gene set enrichment analysis were used across both omics’ layers, generating evidence for the key effects from both differentially expressed genes (DEGs) and more subtle pathway level shifts. Cross-omics correlation analysis was employed to identify how these two omics layers were affected by IVH and by treatment with decorin. Furthermore, little research has been performed on the molecular drivers of the immediate, short-term effects of IVH in the window prior to PHH development. For this reason, leading edge analysis and transcription factor inference were performed to identify the drivers of, and potentially novel targets for, these pathogenic mechanisms.

## 3. Methods

### 3.1. Experimental animals and sample collection

C57BL/6 wild-type mice were obtained from The Jackson Laboratory (Bar Harbor, ME, USA) and bred in the Virginia Commonwealth University School of Medicine at 22°C with a 12:12 light/dark cycle and standard food and water available ad libitum. The design of the experiments, housing, handling, care, and processing of the animals was conducted in accordance with the Guide for the Care and Use of Laboratory Animals and the Animal Welfare Act and the ARRIVE guidelines (Animal Research: Reporting of In Vivo Experiments). All experimental procedures were approved by the Virginia Commonwealth University (AD10003619) Institutional Animal Care and Use Committee.

Donor post-natal day four (PND4) pups were humanely euthanized by decapitation followed by blood collection and lysis in liquid nitrogen for 5 minutes (N = 10). Experimental animals were then anaesthetised followed by bilateral intra-cerebroventricular injection with 6 µl sterile artificial CSF ([aCSF, control] #3525, Tocris Bioscience, Bristol, UK,) or 5 µl blood + 1 µl aCSF (PHH control) / 1 µl decorin (5 mg/ml [#143-DE-100, BioTechne, UK] in distilled water [#46-000-CM, Corning, USA]) (PHH + decorin) per ventricle (Garcia-Bonilla et al., 2026) (Figure 1). Manual intraventricular injections were performed with a 26-gauge needle on a 10 µl syringe (Hamilton Syringe Cat#75N, Hamilton, Reno, NV, USA) with entry 0.8 mm lateral, 1.7 mm anterior, and 1.5 mm deep from the posterior fontanel) and directed to the midline between the orbit level and the ear 1.5 mm deep to the skull surface following our published methods (Garcia-Bonilla et al., 2026). The total volume was injected for 30 seconds into each lateral ventricle. The needle was left for an additional 10 seconds before removal, and mice were allowed to recover from anesthesia and monitored post-induction. Twenty four hours later, animals were anaesthetised with isoflurane (induced at 3% and maintained at 1.5 - 2%, 1 L of oxygen / minute) and underwent MRI scans in a Bruker BioSpec 7T (Bruker) using a T2-weighted TurboRARE sequence (TE = 33 ms, TR = 3000 ms, NEX = 4, 1 repetition, 11 ms echo spacing, rare factor = 8, axial slice orientation, left-right read orientation, 0.5 mm slice thickness, 15 x 15 mm² field of view, 256 x 256 matrix, 0.1 mm slice gap; nominal scan time 6m24s). Animals were then humanely euthanized by transcardiac perfusion with 1 ml ice cold PBS while under deep anaesthesia. Brains were split into two hemispheres, with one immediately dissected and snap-frozen on dry ice, and the other fixed by immersion in 4% paraformaldehyde (#11357638, Fisher Scientific, UK) in PBS overnight at 4°C. All results (transcriptomics, proteomics, and immunostaining) were derived from the same cohort of animals, with animals randomly designated to treatment groups and endpoint analysis technique. Transcriptomic (N = 5 aCSF control, N = 10 IVH, and N = 10 IVH + decorin) and immunostaining (N = 5 per group) was performed on opposite hemispheres each from the same brain. Proteomic analysis (N = 3 per group) was performed on different animals within the same cohort as used for transcriptomics and immunostaining. A total of 45 animals were used in this research, including experimental animals and littermate blood donors.

**Figure 1.**
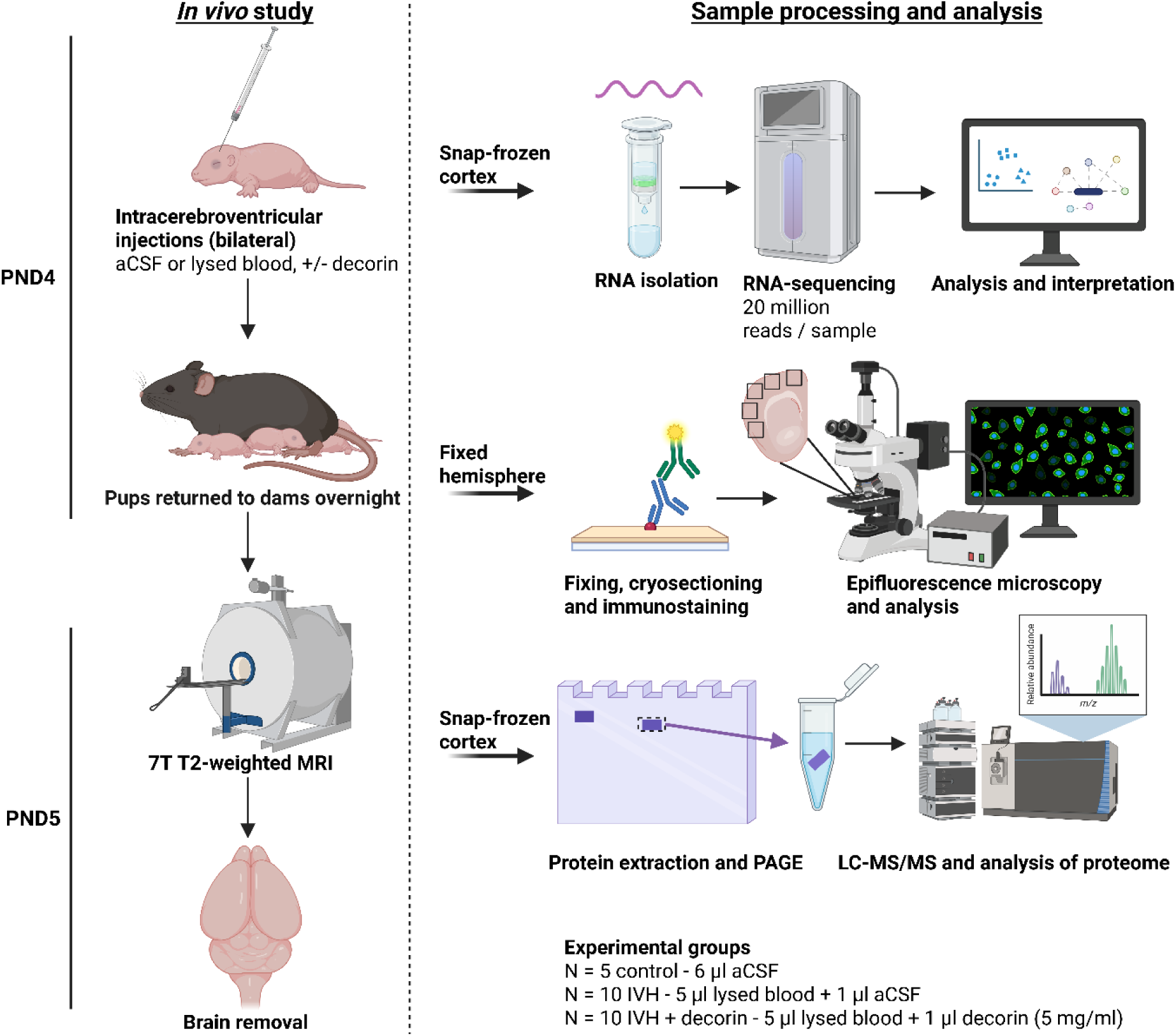
Experimental design and sample processing schematic for RNA-sequencing, immunostaining and proteomics. On PND4, mouse pups received bilateral intracerebroventricular injections of 6 µl artificial cerebrospinal fluid (aCSF control), 6 µl lysed blood from littermate donors (IVH) or 5 µl lysed blood + 1 µl 5 mg/ml decorin (IVH + decorin). At PND5, 7 Tesla T2-weighted MRI was performed, followed by humane culling, perfusion with PBS and brain removal, with each brain split into two hemispheres. One hemisphere had cortical tissue removed and snap-frozen on dry ice. Tissue destined for RNA-sequencing had total mRNA extracted, followed by library preparation and sequencing. Differential gene expression and pathway analyses were then performed. Tissues for proteomic analysis were lysed, reduced, alkylated and trypsinised for LC-MS/MS analysis. The contralateral hemisphere was fixed and processed for immunostaining by cryosectioning.

### 3.2. RNA extraction and sequencing

Snap-frozen cortex tissue from a single hemisphere underwent RNA isolation using a RNeasy micro kit (#74004, Qiagen) following manufacturer’s instructions with a 20 µl elution volume. RNA was then quantified by spectrophotometry (NanoDrop, ThermoFisher). Sample quality control (RIN and concentration) and paired-end mRNA sequencing was performed by a contracted third party (Novogene, UK) using a NovaSeq X Plus (Illumina) at a read depth of 20 million reads per sample.

Alignment was performed against a murine genome (GRCm39) followed by mapping to the same reference genome using the featureCounts tool (Liao et al., 2014) on the Galaxy platform (Abueg et al., 2024) and by quality control using MultiQC (Ewels et al., 2016). Analysis of differentially expressed genes (DEGs) was performed with DESeq2 (Love et al., 2014), generating DEG lists and variance-stabilised transformation (VST) expression scores. DEGs were defined as those with an adjusted p-value (*pAdj)* < 0.1 due to the exploratory nature of this work.

Gene ontology (GO) analysis was performed on all DEGs using goseq (Young et al., 2010). Protein-protein interaction (PPI) analysis was performed using Cytoscape 3.10 with STRING (Szklarczyk et al., 2023), while transcription factor inference was performed using iRegulon (Janky et al., 2014). Over-representation analysis (ORA) was performed on DEGs using the Kyoto encyclopaedia of Genes and Genomes (KEGG) methodology (Kanehisa et al., 2025) using WebGestalt (Elizarraras et al., 2024). Visualisation of cross-omics data was performed using GraphPad Prism version 10 and above.

### 3.3. Tissue processing and immunohistochemistry

Fixed brain hemispheres were cryopreserved by incubation in graded steps (10, 20, 30% w/v) of sucrose in PBS for 24 hours each at 4°C. Tissues were then embedded in optimal cutting temperature media ([OCT] 361603E, VWR, UK) and stored at −80°C prior to sectioning using a Bright OTF7000 cryostat (15 µm sections, −20°C chamber temperature, −23°C sample temperature). Sections were then dried by incubation for two hours on a hotplate set at 37°C prior to storage at −20°C.

Sections for immunofluorescence staining were first washed in PBS + 0.3% Tween-20 ([#CHE3852, Scientific Laboratory Supplies, UK], PBST) to remove OCT media. For primary-secondary immunofluorescence staining, two blocking steps were performed, beginning with immersion in 15% standard goat serum (#16210064, Gibco, UK) in PBST + 2% BSA (A9418, Merck, UK) for 30 minutes at room temperature (RT), followed by three two-minute washes in PBST. A mouse-on-mouse (MOM) blocking step was then performed using anti-mouse F(ab) fragments diluted 1:20 in PBST + 2% BSA, with one hour of incubation at room temperature. Three washes were then performed as previously described, followed by incubation overnight (4°C) in primary antibody diluted in PBST + 2% BSA. Sections were then washed three times (five minutes each at RT), followed by incubation in secondary antibody diluted as previously described for one hour at RT in the dark. Three five-minute washes were performed prior to mounting in media containing DAPI (P36931, ThermoFisher Scientific, UK) and storage at 4°C.

**Table 1.** Antibody dilutions used for immunofluorescent staining of brain sections.

| Antibody | Manufacturer and catalogue number | Dilution |
| --- | --- | --- |
| Anti-NeuN, Mouse | Cell Signalling Technology<br>#94403 | 1:500 |
| Anti-GPX4, Rabbit | Proteintech #67763-1-Ig | 1:400 |
| Anti-GFAP, Mouse | Sigma Aldrich #G3893 | 1:400 |
| Anti-IBA1, Rabbit | Fujifilm Wako #019-19741 | 1:500 |
| Anti-GAD67 | Cell Signalling Technology<br>#41318 | 1:50 |
| Anti-4-HNE | Merck #AB5605 | 1:200 |
| Anti-mouse IgG AF488 plus conjugated,<br>highly cross-absorbed, Goat | ThermoFisher #A32723 | 1:1000 |
| Anti-mouse IgG AF594 plus conjugated,<br>highly cross-absorbed, Goat | ThermoFisher #A32742 | 1:1000 |
| Anti-rabbit IgG AF488 plus conjugated,<br>highly cross-absorbed, Goat | ThermoFisher #A32731 | 1:1000 |
| Anti-rabbit IgG AF594 plus conjugated,<br>highly cross-absorbed, Goat | ThermoFisher #A32740 | 1:1000 |
| Anti-mouse IgG AF660, highly cross-absorbed, Goat | ThermoFisher #A21055 | 1:1000 |
| Anti-mouse F(ab) fragments | Abcam #Ab6668 | 1:20 |

### 3.4. Imaging and image analysis

Epifluorescence images were acquired using a AxioImager Z.1 microscope equipped with Axiocam 305 colour (operating in monochrome mode) and Viluma 7 light source (all Zeiss, Germany) with Zeiss filter sets 96 HE (DAPI), 38 HE (AF488), 50 (AF660) or 91 HE (AF594), maintaining the same imaging settings across all slides within each analysis group. Image processing was performed using FIJI (Schindelin et al., 2012). No image enhancement was performed on images prior to analysis unless specified, and any changes to representative images (background subtraction and modification of window and level) were performed uniformly across a comparison group. Five regions of interest (ROIs) were captured from each cortex, utilising the DAPI channel to avoid bias. Automated analysis was performed in QuPath (Bankhead et al., 2017), segmenting cells using DAPI^+^ or NeuN^+^ cell detection or trained object classifiers, followed by analysis of cell positivity over a pre-set threshold or extraction of mean fluorophore intensity per cell. Values obtained from five ROIs were then averaged to give a mean value for each animal.

### 3.5. Protein extraction, proteomics and cross-omics analysis

Samples were lysed in 250 µl radioimmunoprecipitation assay (RIPA) buffer supplemented with protease (#11697498001, Roche, Switzeland) and phosphatase inhibitors (#4906845001, Roche, Switzerland) per manufacturer’s instructions. Samples were then incubated on a rocker at 4°C for 2 h and centrifuged at 10,000 × *g* for 20 min at 4°C. Supernatant was harvested, aliquoted and stored at −80°C until required. Protein concentration was assessed by BCA assay (A55864, ThermoFisher Scientific, UK), following the manufacturer’s instructions.

5-10 μg of lysate was separated in 8 mm on mPAGE 1 mm 12% Bis-Tris 10 or 12 well gels (MP12W10 / MP12W12, Merck, UK) prior to Coomassie Blue staining (GEN-QC-STAIN-1L, Generon). Bands were cut and destained with extraction buffer (50% acetonitrile, 100 mM ammonium bicarbonate, 5 mM DTT, 16 h, 4 °C). Samples were then alkylated (40 mM chloroacetamide, 160 mM ammonium bicarbonate, 10 mM TCEP, 20 min, 70 °C), dehydrated in 100% acetonitrile, and air-dried at 37 °C, followed by overnight digestion with sequencing grade trypsin (#V5111, Promega, UK).

Peptide digests were separated using a Vanquish Neo UHPLC system using a 50 cm EASY-Spray™ PepMap™ Neo analytical column. Tryptic peptides were eluted over an 80-min linear gradient from 4–36% acetonitrile, 0.1% formic acid at 300 nL/min and electrosprayed at 1,800 V into an Orbitrap Ascend Tribrid mass spectrometer. The instrument was operated in data-dependent acquisition (DDA) mode. Full MS scans were acquired in the Orbitrap at 120,000 resolution with a normalized AGC target of 250%, maximum injection time of 50 ms, and dynamic exclusion of ±10 ppm for 15 s. MS/MS spectra were generated using HCD at 30% collision energy and detected in the ion trap with turbo scan rate, using a 100% AGC target and automatic maximum injection time.

Raw files were processed using MaxQuant 2.1.0 and Perseus 1.6.12. with a recent download of the UniProt *Mus musculus* reference proteome (UP000000589), and the sequence of human decorin (P07585). A decoy dataset of reverse sequences was used to filter false positives. Trypsin digest with 2 missed cleavages and a minimum peptide length of 7 residues was used with both peptide and protein FDR thresholds set to 0.01. Differential abundance calculations between conditions were performed in Perseus, using permutation-based FDR correction, with FDR = 0.05, S0 = 0.1 and 250 randomisations.

Due to the lower coverage provided by proteomic analysis compared with RNA-sequencing, gene set enrichment analysis (GSEA) was utilised in addition to the over-representation analysis used for transcriptomic analyses. This was then compared with transcriptomic data using a cross-omics approach. KEGG and GO GSEA was performed on a ranked list of the detected, filtered proteins and genes separately using WebGestalt. Enrichment scores (normalized enrichment score, NES) and Benjamini-Hochberg corrected false discovery rates (FDR) were calculated for each dataset. An exploratory false discovery rate (FDR) of 0.25 was set for KEGG and GO GSEA, as recommended in the original GSEA framework (Subramanian et al., 2005). Hypothesis-driven proteomic analysis was also performed using predefined protein lists (representing neuronal integrity, microglial biology, and oxidative stress/iron homeostasis). Proteins of interest, including NEFL and NEFM, were used to inform the generation of these protein lists, with the rationale for their selection described in Supplemental Table S2. Protein abundance was assessed using log₂ LFQ intensities and visualised using heatmaps and individual protein plots. Within these targeted analyses, proteins were selected based on biological relevance and were not required to meet proteome-wide multiple testing correction thresholds for inclusion.

Correlation analysis of the alignment between transcriptomic and proteomic data was performed by extracting all GO terms for each ontology (biological process, cellular component, molecular function) with FDR < 0.25. Cross-omics (transcript / protein) analysis was then performed by examining agreement between detected gene-protein pairs and ontologies across both omics’ layers.

Terms present in both GO datasets were then analysed further with leading edge analysis, where leading edge subsets were extracted to identify genes or protein which most strongly contributed to enrichment of the term. Leading edge occurrence scores were generated for each gene or protein as the number of GO terms in which it appeared within the selected subset. These leading edges were identified as contributors to pathways enrichment rather than as strictly differentially expressed genes or proteins.

### 3.6. Statistical analysis

All statistical analysis was performed in GraphPad Prism unless otherwise stated. Statistical tests are described in the relevant figure legends. Biological replicates are defined as separate animals. Investigators were not masked to the identity of animals whilst performing experiments *in vivo*, however data analysis was performed on anonymised files where possible, with filenames hidden during image quantification.

## 4. Results

### 4.1. Neonatal IVH significantly modulates injury markers in the cortical transcriptome

Post-natal day 5 (PND5) animals were first assessed for ventriculomegaly using MRI, 24 hours after bilateral intracerebroventricular injection with aCSF, blood + vehicle (IVH) or blood + decorin (IVH + decorin). Quantification of combined lateral ventricle volume showed no statistically significant differences in ventricular volume at this timepoint (Figure S1).

Quality control of 25 RNA samples (N = 5 aCSF control [group A], N = 10 IVH [group B], N = 10 IVH + decorin [group C]) showed that each extracted RNA sample had a high concentration and purity, as assessed by Nanodrop spectrophotometry (Table S1). Following RNA-sequencing, mapping quality control showed that each sample has a minimum of 20 million unique reads assigned (Figure S2A), and a mean GC content of ∼50% (Figure S2B). Principal component analysis (PCA, Figure S2C) and hierarchical clustering of the top 500 differentially expressed genes (DEGs, Figure S2D) revealed distinct separation of the transcriptomic profile of aCSF control animals from those subjected to IVH and IVH + decorin, with the exception of sample C4, though this was not excluded from subsequent analyses to ensure that biological variability was captured.

Analysis of differentially expressed genes (DEGs, defined as pAdj < 0.1 due to the exploratory nature of this work) revealed a total of 196 DEGs (87 upregulated and 109 downregulated) when comparing IVH animals to aCSF control, shown by volcano plot with example DEGs labelled in Figure 2A. Most log_2_(fold change [FC]) values were below 0.5, indicating a relatively small magnitude of change in gene expression across these groups.

**Figure 2.**
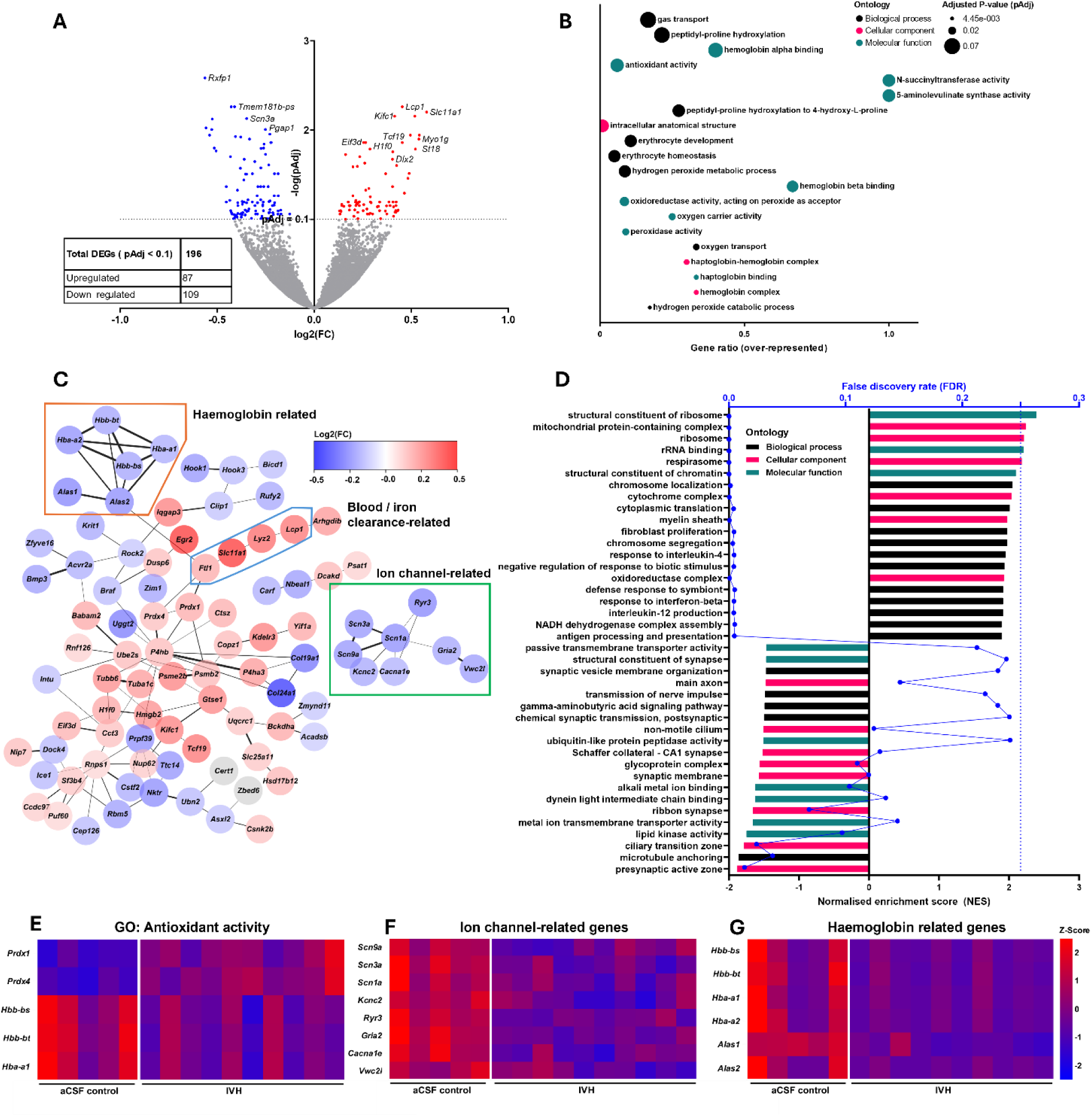
Transcriptomic data enrichment reveals enrichment of genes related to antioxidant activity, ion channels and blood clearance. A) IVH vs aCSF volcano plot with representative DEGs labelled and inset table summarising number of DEGs and split up and down regulated DEGs. Upregulated genes (shown in red) in this comparison are defined as those with significantly greater expression in IVH animals than aCSF control. B) Gene ontology (GO) of top 20 over-represented terms between aCSF control and IVH brains, showing gene ratio (proportion of genes within GO term which are differentially expressed in this dataset), GO, adjusted P-value and ontology (black = biological process, pink = cellular component, teal = molecular function). C) STRING-enrichment of protein-protein interactions within DEGs (IVH compared to aCSF control), shown as nodes with non-connected genes excluded. Colour depicts Log_2_(FC) (red = upregulated, blue = downregulated), thickness of interconnecting lines depicts confidence of protein-protein interaction (thicker = higher confidence). D) GSEA analysis of transcriptomic data, identifying gene ontologies affected by IVH with enrichment of “respirasome”, “response to interleukin 4” and “oxidoreductase complex”. Depleted ontologies included “presynaptic active zone”, “synaptic membrane” and “transmission of nerve impulses”. E) Heatmap of DEGs within the GO term Antioxidant activity (molecular function, GO:0016209). F,G) Heatmap of protein-protein interaction (PPI) sub-network related to ion channel gene expression (E, outlined in green in C) and haemoglobin related genes (F, outlined in orange). DEGs defined as pAdj < 0.1.

Gene ontology (GO) over-representation analysis (ORA), which only includes DEGs within the analysis, was then performed. GO analysis of IVH relative to aCSF control animals revealed significant over representation of blood-related GO terms (Figure 2B), including “hemoglobin alpha binding”, “hemoglobin beta binding” and “erythrocyte homeostasis”. Terms linked to reactive oxygen species (ROS) defence, including “antioxidant activity”, “peroxidase activity”, “oxygen transport” and “oxygen carrier activity” were also over-represented (Figure 2B).

Mapping of DEGs onto a protein-protein interaction (PPI) network (Figure 2C) showed significant downregulation of a cluster of blood related genes (*Hbb-bt*, *Hbb-bs*, *Hba-a1*, *Hba-a2*, *Alas1* and *Alas2)* following IVH. A group of genes (*Ftl1*, *Slc11a1*, *Lyz2* and *Lcp1*) related to the response of tissue to blood and iron showed significant upregulation (Figure 2C). Additionally, a cluster of ion-channel related genes, including *Scn3a*, *Scn9a*, *Scn1a*, *Kcnc2* and *Cacna1e,* were significantly downregulated in IVH animals.

Broader, higher coverage GO GSEA including all genes was then performed (Figure 2D). Here, significant enrichment was seen in ontologies relating to inflammation including “response to interleukin-4, “response to interferon-beta” and “interleukin 12 production”, whilst significant depletion was noted for ontologies relating to neurons and synapses including “presynaptic active zone”, “metal ion transmembrane transporter activity”, “main axon” and “chemical synaptic transmission, postsynaptic”.

The GO term “antioxidant activity” was also represented by a heatmap of significantly over-represented genes within this term, revealing IVH-induced upregulation of *Prdx1* and *Prdx4* and downregulation of *Hbb-bs*, *Hbb-bt* and *Hba-a1* (Figure 2E). Ion-channel and haemoglobin-related genes, identified through PPI analysis, were also plotted in a similar fashion (Figure 2F and G respectively).

### 4.2. Cross-omics analysis reveals convergent neuronal depletion alongside ferroptosis and phagocytosis enrichment in neonatal intraventricular haemorrhage

Following initial transcriptomic analysis, we performed proteomic analysis to identify proteins altered following IVH. Volcano plot representation of proteomic data indicated significant downregulation of proteins including NEFM, NELF, and CACNG8 and significant upregulation of FTH1, FTL1, and KNG1 (Figure 3A) in IVH. Proteomic GSEA analysis using the Kyoto encyclopaedia of genes and genomes (KEGG, Figure 3B) revealed significant enrichment of pathways including “Ribosome”, “Ferroptosis”, and “Complement and coagulation cascades”. Significantly depleted pathways included “Cell adhesion molecules”, “Gap junction”, and “Synaptic vesicle cycle”. Consequently, we focussed on proteins related to ferroptosis, iron handling, and antioxidant defence due to implication of these pathways in previous transcriptomic analysis. The heatmap shows a trend towards upregulation of these proteins following IVH (Figure 3C). Within the ferroptosis and iron handling protein panel, FTH1, FTL1, and ACOX1 showed a significant increase in abundance following IVH (P < 0.0001, P < 0.000 and P = 0.0229, respectively), while only PRDX1 was significantly raised (P = 0.0278) within the antioxidant related protein panel (Figure 3D). Immunostaining was then employed to study GPX4, a key antioxidant involved in ferroptosis defence that was not detectable during our proteomics analysis, and 4-hydroxynonenal (4-HNE), a chemical marker of lipid peroxidation. GPX4 was significantly reduced in IVH brains compared with that of aCSF control animals (P = 0.0247, Figure 3E-F) however neuronal positivity (in approximately cortical layers II through III) for 4-HNE did not show any significant changes (Figure G-H).

**Figure 3.**
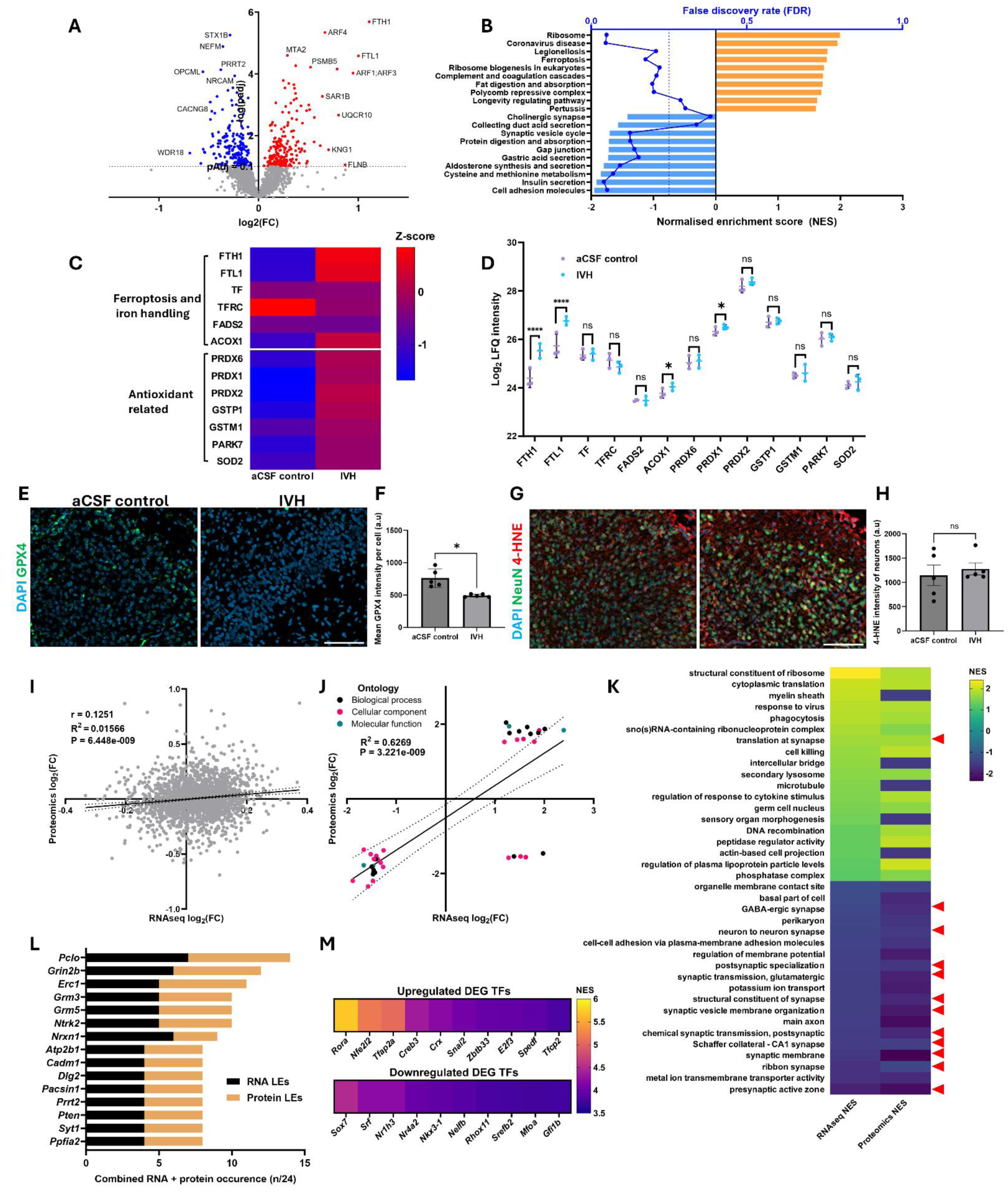
Cross-omics integration reveals convergent molecular pathways in neonatal intraventricular haemorrhage. A) Volcano plot representation of IVH proteome, with representative proteins labelled. B) Gene set enrichment analysis (GSEA) of proteomic data revealed key KEGG pathways enriched by NIVH (orange bars), including “Ferroptosis” and “Complement and coagulation cascades”, whilst depleted pathways (blue bars) included “Cell adhesion molecules”, “Gap junction”, and “Synaptic vesicle cycle”. Benjamini-Hochberg corrected false discovery rate (FDR) is also presented along the top X-axis in blue. C) Curated gene set enrichment of proteins related to ferroptosis, iron handling and antioxidant defence depicted by heatmap and by protein abundance using LFQ values (D). Statistical analysis of protein abundance was performed using t-test in Perseus with permutation-based FDR correction (FDR = 0.05, S0 = 0.1, 250 randomisations. N = 3 biological replicates. E-F) Immunofluorescence staining and image analysis of glutathione peroxidase (GPX4) and neuronal expression of 4-HNE (G-H). Statistical analysis of GPX4 immunofluorescence imaging performed using one-way ANOVA with Tukey’s test. Neuronal 4-HNE positivity analysed using Kruskal-Wallis test due to failing normality testing. All data are mean ±SD. I) Cross-omics concordance was assessed by correlation of all 2137 matched genes and proteins using Pearson’s *r* test. J) Correlation between gene ontologies was then performed, again using Pearson’s *r* test. 95% confidence intervals depicted on panels D and E using dotted lines. K) Heatmap demonstrating the relative magnitude and direction of enrichment between transcriptomic and proteomic gene GSEA ontology analysis. Enriched terms (yellow) across all datasets included “cell killing” and “phagocytosis”, with depletion (purple) noted in multiple terms related to the synapse and neurons (highlighted with red triangles). α = 0.25 for all GSEA. L) Leading edge genes (black) and proteins (brown) forming leading edges for synapse-related GO terms (highlighted by red triangles in K), represented as a proportion of 24 combined occurrences (gene and protein) across 12 GO terms. M) Inferred transcription factor (TF) activity from up-and down-regulated DEGS in IVH compared with aCSF control animals.

To determine whether these pathway-level alterations were consistent between omics layers, we next performed cross-omics correlation analysis. Pearson’s *r* correlation was utilised on all detected gene-protein pairs (n = 2137), identifying a weak but statistically significant positive association between these omics’ layers (*r* = 0.125, P = 6.45×10^-9^, Figure 3I). We next examined the concordance between transcriptomic and proteomic pathways by plotting all GSEA gene ontology pairs with FDR <0.25 in both layers (Figure 3J, n = 38), with Pearson’s *r* test showing a strong correlation between transcriptomic and proteomic enrichment scores (R^2^ = 0.627, P = 3.22×10^-9^). All paired gene ontologies with FDR < 0.25 were then plotted by heatmap to show their relative magnitude and direction of enrichment (Figure 3K). This cross-omics analysis identified concurring high enrichment of ontologies including “phagocytosis”, “cell killing”, and “regulation of response to cytokine stimulus”. Significant depletion was seen in ontologies including “metal ion transmembrane transporter activity”, “chemical synaptic transmission, postsynaptic”, “regulation of membrane potential”, and “cell to cell adhesion via plasma-membrane adhesion molecules”. All paired ontologies which displayed depletion in the transcript layer showed matching depletion in the protein layer (n = 19, 100% concordant), while 5 of 19 ontologies showing transcript enrichment had a discordant depletion at the protein level (73.7% concordant).

Notably, 12 of 38 pairs were synapse-related GO terms (highlighted in Figure 3K with red triangles); therefore, leading-edge (LE) analysis was performed on these subsets from both transcriptomic and proteomic GSEA analyses to identify which genes contributed to this enriched signal. The top 15 overall LEs (ranked by summed RNA + protein occurrence) were then depicted in Figure 3L. The top-ranking LE was *Pclo*, occurring in 7/12 transcriptomic GO terms and 7/12 proteomic GO terms, followed by *Grin2b* (6/12 and 6/12, transcriptomic and proteomic) and *Erc1* (5/12 and 6/12, respectively). While these LE genes may not be differentially expressed in isolation, they consistently contribute to synaptic pathway depletion across multiple GO terms at transcript and protein level.

To identify possible drivers of the effects of IVH, transcription factor (TF) inference was then performed on DEGs obtained from RNA-sequencing data. TF inference with iRegulon (Figure 3M) suggested *Rora*, *Nfe2l2*, and *Tfap2a* (NES 5.714, 5.172 and 5.065 respectively) as the top 3 putative TFs due to enrichment of their binding motifs amongst the upregulated DEGs. TF inference of downregulated DEGs suggested enrichment of motifs for *Sox7*, *Srf*, and *Nr1h3*, though NES values for these were generally lower than those inferred from the upregulated DEG dataset (NES = 4.501, 4.234, and 4.223 respectively). Overall, both transcriptomics and proteomics data suggest IVH increases inflammation, phagocytosis, haemoglobin and iron clearance, and reduces ion channel abundance and synaptic/neuronal pathways within the cortex.

### 4.3. Transcriptomics suggests decorin modulates inflammation, microglial activation and phagocytosis after IVH

ORA was then applied to transcriptomic data comparing decorin-treated IVH with untreated IVH to identify high-magnitude changes driven by differential gene expression. 54 DEGs (pAdj < 0.1) were identified, wherein 54 genes were upregulated and 1 gene was downregulated (Figure 4A). A directionality plot (**Error! Reference source not found.**B), was then used to compare log_2_(FC) values of IVH-induced DEGs with corresponding log_2_ fold changes of the same genes after decorin treatment. Whilst these decorin-induced changes were not necessarily statistically significant, this provided an overall indication of the directionality of transcriptomic changes and candidate genes for further analysis. Several genes were significantly upregulated by IVH and then further upregulated by decorin (suggesting decorin-induced enhancement, Figure 4B top right quadrant), including *Myo1g*, *Slc11a1*, *Lcp1*, and *Oas1a*. Rescued genes (upregulated by IVH and downregulated by decorin; Figure 4B bottom-right quadrant) included *St18*, *Deup1*, *Dlx2*, and *Tcf19*. A large number of genes were rescued in the opposite direction (downregulated by IVH and upregulated by decorin, Figure 4B top left quadrant), including *Col24a1*, *Mtcp1*, and *Rxfp1*. A limited number of genes were downregulated by IVH and then further downregulated by decorin (Figure 4B, bottom left quadrant).

**Figure 4.**
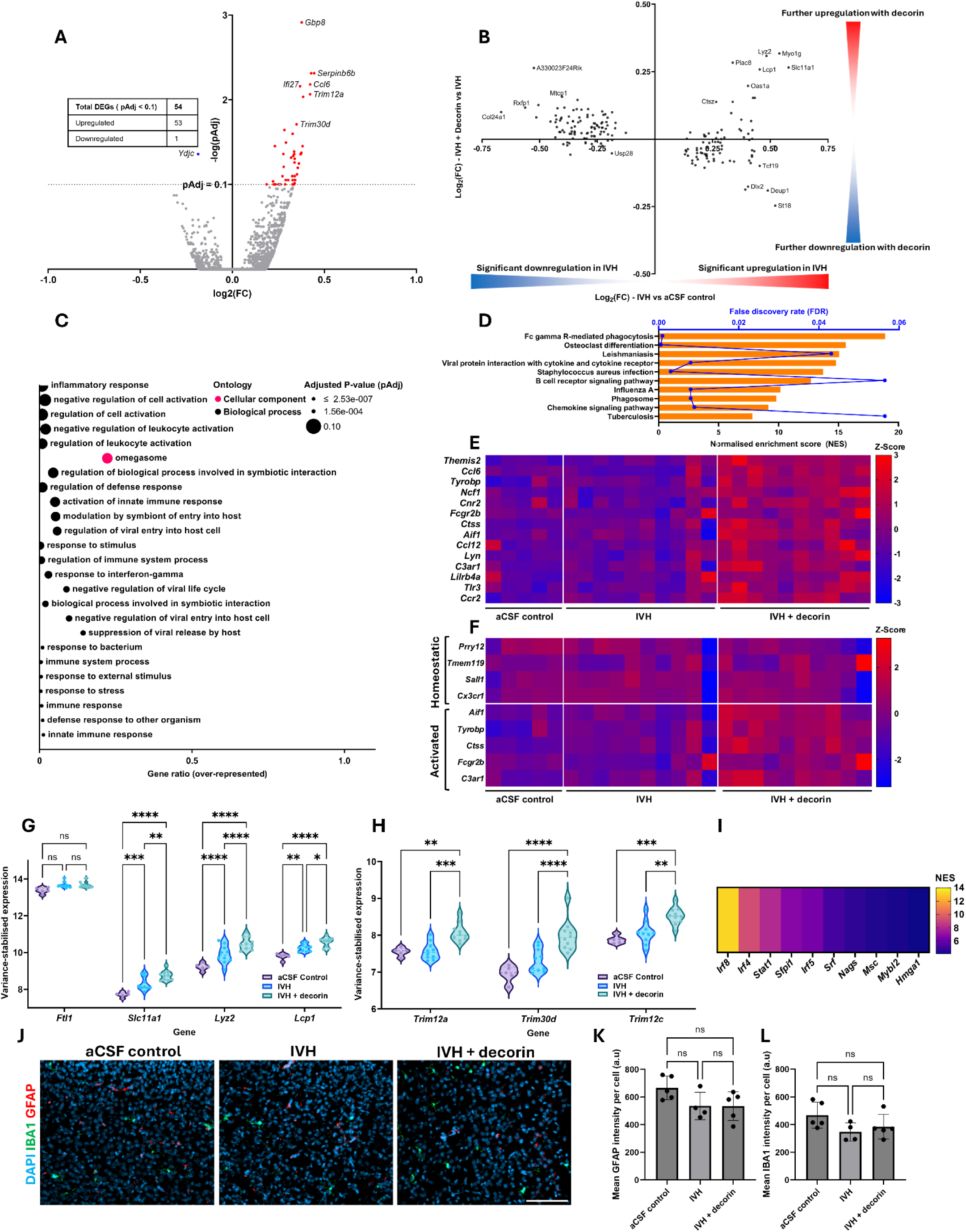
GO, KEGG pathway and DEG transcriptomic analysis for decorin-treated IVH. A) Volcano plot of IVH + decorin vs IVH contrast with representative DEGs labelled and inset table summarising number of DEGs and split up and down regulated DEGs. Upregulated genes (shown in red) in this comparison are defined as those with significantly greater expression in IVH + decorin animals than IVH only animals. B) Directionality plot showing all DEGs from IVH vs aCSF comparison, plotted by log2FC along the X-axis. Y-axis depicts the corresponding log2FC for the same genes in IVH + decorin vs IVH, showing how decorin modulated IVH-induced DEGs. DEGs defined as pAdj < 0.1. C) GO for over-represented terms from IVH + decorin DEGs, showing gene ratio (proportion of genes within GO term which are differentially expressed in this dataset), GO name, adjusted P-value, and ontology (green = biological process, red = cellular component). D) KEGG ORA of DEGs, with orange depicting positively enriched terms plotted by normalised enrichment score (NES) on the bottom X-axis in black. Benjamini-Hochberg corrected false discovery rate (FDR) is also presented along the top X-axis in blue. E) Heatmap of DEGs from GO term “Inflammatory response”. F) Heatmap of curated microglial geneset using a gene set enrichment analysis approach. G, H) Variance stabilised expression plots of DeSeq2 DEGs from blood iron related PPI clusters (Figure 2C) and the selected GO term “Omegasome”. Statistical analysis performed by two-way ANOVA with Šídák’s multiple comparisons test. I) Inferred transcription factor (TF) activity from down-regulated DEGS in IVH+decorin compared with IVH animals. J-L) Immunofluorescence detection and quantification of astrocytes (GFAP) and microglia (IBA1). Statistical analysis of immunofluorescence imaging performed using one-way ANOVA with Tukey’s test, one missing replicate in IVH group of panels K and L due to loss of tissue from slide, mean ±SD. * *P* < 0.05, ** *P* < 0.01, *** *P* < 0.001, **** *P* < 0.0001, α = 0.05.

ORA GO and KEGG enrichment was then performed to identify any ontologies driven by decorin-mediated differential gene expression. Decorin treatment appeared to have a significant impact on inflammation, with over-representation of GO terms including “inflammatory response”, “negative regulation of leukocyte activation”, and “regulation of defense response” (Figure 4C). KEGG ORA was also performed (Figure 4D), with “Fc gamma R-mediated phagocytosis” being the most over-represented, significantly enriched KEGG term, followed by “Phagosome”. One of the genes highlighted by the heatmap of over-represented genes in the “inflammatory response” GO term (Figure 4E) was *Aif1*, which is a common marker of microglia within the CNS. Microglia are the resident inflammatory cells within the brain and are known to play key roles in the response of the brain to injury and disease. Therefore, we focussed on genes related to microglial homeostasis and activation to identify changes in microglia with decorin treatment. Expression of homeostatic genes *P2ry12*, *Tmem119*, *Sall1*, and *Cx3cr1* was broadly consistent across all conditions, whilst genes associated with microglial activation and immune signalling (*Aif1*, *Tyrobp*, *Ctss*, *Fcgr2b*, *C3ar1*) showed a trend towards upregulation in decorin-treated IVH compared to IVH and control animals (Figure 4F).

Microglia are thought to contribute to the brain’s immune response to blood breakdown products following IVH (Umpornpun et al., 2026). As IVH had induced the modulation of genes related to blood and iron clearance through phagocytosis linked pathways (Figure 2), these genes were analysed here to identify whether decorin modified their expression further. Plotting of VST-normalised expression (Figure 4G) revealed that *Slc11a1*, *Lyz2*, and *Lcp1* showed significant stepwise increases in expression from aCSF control to IVH to IVH + decorin groups. In addition to KEGG analysis showing phagocytosis as enhanced with decorin treatment, GO analysis showed over-representation of the “omegasome” term (Figure 4C), which has a role in autophagy (Nähse et al., 2024). Three omegasome-linked genes (*Trim12a*, *Trim30d*, and *Trim12c*) were shown to have no significant difference in VST-normalised counts between aCSF control and IVH (P > 0.05). However, significant differences were noted between IVH and IVH + decorin groups, with VST normalised counts for *Trim12a* increasing from 7.571 to 8.070 (P = 0.0006), *Trim30d* from 7.317 to 7.986 (P < 0.0001), and *Trim12c* from 8.065 to 8.524 (P = 0.0017) with decorin treatment (Figure 4H). Finally, TF inference of upregulated DEGs suggested *Irf8*, *Irf4*, and *Stat1* as the top three active transcription factors (NES = 13.19, 9.29 and 8.07 respectively, Figure 4I), of which *Irf8* is highly expressed in postnatal microglia.

To establish whether the increase in inflammatory genes within the cortex was due to an increase in microglia / macrophages or astrocyte cell number we performed immunofluorescence staining of the cortex with an antibody to IBA1, quantified through approximately cortical layers II-III (Figure 4J-L). No significant changes were noted following IVH and decorin treatment (P > 0.05). Overall, these data suggest that decorin may promote a shift to a more responsive or activated microglial phenotype within the cortex to boost the clearance of blood products.

### 4.4. Decorin reduces neuronal loss in the cortex following IVH

Following this transcriptomic ORA approach, protein expression was then assessed between IVH and IVH + decorin. This approach identified 677 differentially expressed proteins using a relaxed *pAdj* < 0.1 cutoff (Figure 5A, 335 upregulated and 342 downregulated) and an upregulation of proteins including NEFM and NELF, two neuronal filament proteins. Proteomic GSEA using KEGG (Figure 5B) showed that decorin induced significant enrichment of terms including “Phagosome” and “Proteasome”. Significant depletion was only noted for the terms “Spliceosome” and “Ribosome”. Thus, these analyses supported some of the transcriptional changes associated with phagocytosis.

**Figure 5.**
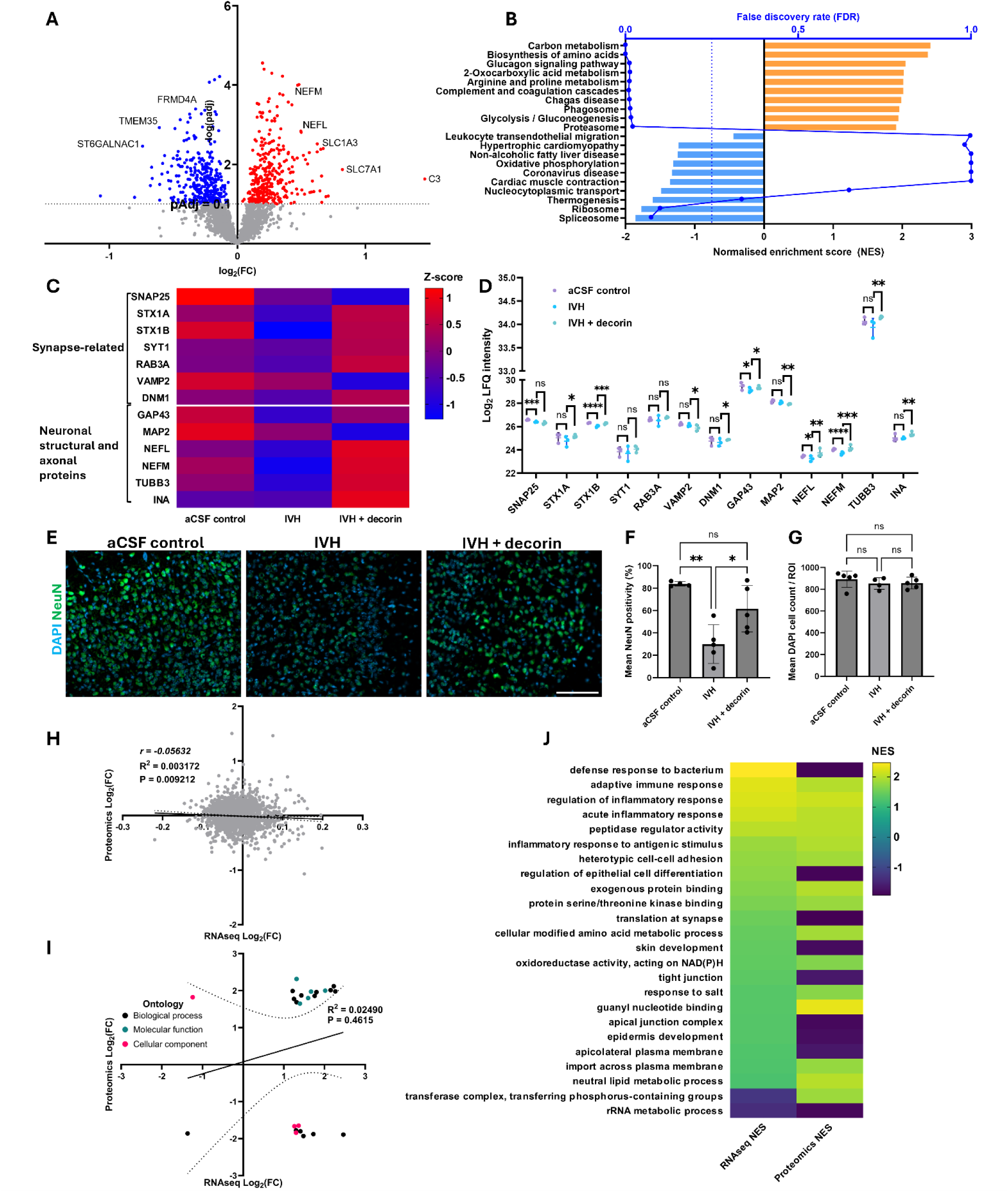
Decorin alters transcript–protein relationships in neonatal IVH. A) Protein volcano plot, with selected proteins labelled. B) Proteomic GSEA using KEGG showed enrichment of terms including “Carbon metabolism”, “Phagosome” and “Proteasome”, with depletion of “Spliceosome” and “Ribosome” (orange = enriched, blue = depleted). Benjamini-Hochberg corrected false discovery rate (FDR) is also presented along the top X-axis in blue. C) Heatmap representation of the proteomic expression of selected gene sets relating to synapse functionality and neuronal structure, in addition to protein abundance utilising log2 LFQ intensity values (D). E) Immunofluorescence staining and image analysis (F) of cortical neurons (NeuN) and of total DAPI+ve cells in the cortex (G). One replicate excluded in aCSF control column of NeuN (C) due to failing ROUT outlier test with Q = 1%, one missing replicate in IVH group in panel J due to loss of tissue on slide – all others N = 5. Data with missing or excluded replicates were analysed with one-way ANOVA with Holm-Šídák’s multiple comparisons test. DAPI cell count analysed using Kruskal-Wallis test due to failing normality testing. H) Cross-omics correlation of all 2141 gene-protein pairs, assessed using Pearson’s *r*, indicating a decoupling of these two omics layers. I) Assessment of correlation between gene ontologies significantly modulated in both omics layers (n = 24, FDR < 0.25) using Pearson’s *r* test. 95% confidence intervals depicted on panels H and I using dotted lines. J) Heatmap demonstrating the relative magnitude and direction of enrichment between transcriptomic and proteomic gene GSEA ontology analysis within the IVH + decorin vs IVH contrast. Enriched terms (yellow) across all datasets included “adaptive inflammatory response” and “regulation of inflammatory response”, with depletion (purple) only noted across both layers for “rRNA metabolic process”. All data are mean ±SD.

As neuronal markers appeared to be elevated with decorin treatment, we performed expression analysis on a set of key proteins implicated in both synapse function and neuronal/axonal structure, including STX1A/B, NEFL, NEFM, and INA, which is described in supplemental Table S2 and derived from literature. Heatmap representation of protein Z-scores revealed a trend of reduced abundance of these proteins in IVH animals, followed by a return towards the control baseline in those treated with decorin (Figure 5C). This was further analysed through plotting log_2_ LFQ intensities of these proteins, revealing that decorin successfully rescued the IVH-induced reduction in STX1B, GAP43, NEFL, and NEFM (Figure 5D). These proteins were all significantly reduced by IVH (P < 0.0001, = 0.0263, = 0.0181 and < 0.0001 respectively) and then significantly increased from this level in IVH + decorin animals (P = 0.0003, = 0.0184, = 0.0015, = 0.0001). The expression of two other neuronal proteins (SYT1, RAB3A) was not significantly altered in any tested condition (P > 0.05 for all). SNAP25 expression was significantly reduced after IVH but not significantly altered by decorin treatment (P = 0.0006 and > 0.05). STX1A, DNM1, TUBB3, and INA were not significantly altered by IVH (P > 0.05), but were all increased in IVH + decorin brains compared to IVH alone (P = 0.0115, 0.0304, 0.0029, and 0.0002). VAMP2 was not significantly different between control and IVH groups but was downregulated in IVH + decorin animals (P = 0.0219).

This was further supported by immunostaining data, with cortical NeuN cell positivity falling from 83.65% in control brains to 29.89% in IVH (P = 0.0017, Figure 5E and F). Decorin treatment significantly restored NeuN cell positivity to 61.48% (P = 0. 0214) which was not statistically significant from control brains (P > 0.05). A similar trend was observed when cortical NeuN staining in layers II-III was analysed as mean intensity per cell (Supplementary Figure S3). However, the GABAergic-specific neuron marker GAD67, utilised due to cross-omic depletion of GABAergic synapse terms with IVH, showed no significant difference between any tested condition (P > 0.05 for all, Supplementary Figure S4). No significant difference was noted in total DAPI^+^ cells within the cortex (Figure 5G). This data suggests that decorin treatment acts to protect the cortical neurons and prevent neuronal loss 24 hours following IVH.

### 4.5. Decorin treatment caused a loss of transcriptomic and proteomic co-ordination

Gene-protein correlation (Figure 5H) between IVH and IVH + decorin was shown to be very weak and slightly negative (*r* = −0.05632, R^2^ = 0.003172) but statistically significant (*P* = 0.009212, likely driven by a high sample size of n = 2141 gene-protein pairs), indicating minimal overall agreement between gene expression and protein levels. This was confirmed following correlation analysis of gene ontologies (Figure 5I), which showed minimal, non-significant agreement between omics layers (R^2^ = 0.0249, *P* = 0.4615), suggesting a loss of the coordinated pathway level responses seen with IVH (Figure 3). GO GSEA was then performed on both omics layers, and any with FDR < 0.25 in both were plotted by heatmap (Figure 5J). Marked, concordant enrichment was noted for several terms related to the inflammatory response, including “regulation of inflammatory response” and “acute inflammatory response”. Concordant depletion was only observed for “rRNA metabolic process”. This suggests decorin treatment has uncoupled the relationship between transcript and protein and is modulating proteins independently of their mRNA transcripts in IVH.

## 5. Discussion

We have described a cross-omics study of neonatal IVH with a decorin treatment arm, using transcriptomics, proteomics, and immunostaining to identify cortex-specific responses 24 hours after injury. We also identified there was not a significant difference in ventricular volume at this timepoint, indicating a potential window to prevent the progression of IVH into PHH, one of the most serious complications that can follow IVH.

An integrative, cross-omics approach identified depletion of gene ontologies related to neurons, synapses, and metal iron transport in IVH brains at both the gene and protein levels. These two omics layers were shown to be significantly concordant in IVH, both at the level of individual gene-protein pairs and between ontologies, suggesting a unified biological response to the injury. This was further supported by protein abundance analysis, including analysis using a curated protein set based on existing literature (supplemental materials Table S2) (Benowitz & Routtenberg, 1997; Dehmelt & Halpain, 2005; Dixon & Stockwell, 2018; Ferguson & De Camilli, 2012; Y. Hu et al., 2025; Puri et al., 2023; Shim & Kim, 2013; Südhof, 2004, 2013; Tew & Townsend, 2012; Yamane et al., 2022; Yuan et al., 2017). This revealed reductions in cortical levels of key neurofilaments NEFL and NEFM, neuronal growth protein GAP43 (Benowitz & Routtenberg, 1997), and the synaptic transmission protein STX1B (Südhof, 2004, 2013) after IVH. Dissociation of these neurofilaments (key components of the neuronal cytoskeleton) and subsequent reductions in neuronal stability are hallmarks of traumatic brain injury (Siedler et al., 2014; Yuan et al., 2017). These findings were further reinforced by a reduction in neuronal staining with NeuN in the cortex of IVH mice compared with aCSF controls. Post-mortem studies of human preterm infants have previously shown reductions in the number of intermediate progenitor cells (which ultimately form cortical pyramidal neurons) in IVH brains compared to controls, indicating that this reduction in neurons is a pathological indication of human neonatal IVH (Sharma et al., 2022).

Decorin treatment ameliorated these effects in our neonatal mouse model, restoring the abundance of NEFL, NEFM, GAP43, and STX1B to near control levels, in addition to restoring NeuN positive neuronal staining, suggesting decorin may have some neuroprotective properties in IVH. Furthermore, decorin increased levels of STX1A (a synaptic transmission protein with similar function to STX1B (Kofuji et al., 2014)), TUBB3 (a key microtubule protein involved in axon guidance and maintenance (Puri et al., 2023)) and DNM1 (involved in synaptic vesicle recycling (Ferguson & De Camilli, 2012)) above levels found in IVH brains, even though this was not significantly less than that in control cortex tissue.

Across both omics layers there was an emphasis on the GO term “cell killing”, oxidative stress, and ferroptosis (iron-mediated, non-regulated cell death) pathways. Our data suggests these enrichments could be specific to neurons, as these were the only cell type to show a significant reduction following IVH, though single-cell analysis would be required for validation.

Oxidative stress and ferroptosis are widely known to be induced by tissue damage after haemorrhagic stroke and have been the focus of a multitude of therapeutic studies (Cheng et al., 2026; Dixon et al., 2012; Duan et al., 2016). The effect of ferroptosis on periventricular tissues, including the choroid plexus and periventricular white matter, has been well documented. PHH has been associated with increased expression of ferroptosis markers in the choroid plexus (Meng et al., 2023) and chelation of free iron has been shown to reduce white matter injury and neurological dysfunction (Cheng et al., 2026). However, these prior studies have not examined the possible implication of ferroptosis in cortical tissue damage after IVH.

Here, transcriptomic ORA of cortical tissue revealed that some of the strongest RNA-level responses to IVH was the upregulation of a cluster of linked genes including *Ftl1*, *Slc11a1* and *Lyz2. Ftl1* and *Slc11a1* have been shown to be strongly linked to ferroptosis and iron-clearance (Shi et al., 2024; Zhou et al., 2025). Both FTL1 and its heavy-chain partner FTH1 also showed this same enrichment at the protein level. FTL1 is suggested to act as an iron-storage protein, with accumulation resulting in neurodegeneration through synaptic damage (Dixon & Stockwell, 2018; Remesal et al., 2025). SLC11A1 is a macrophage-specific membrane transporter of iron (Liziczai et al., 2025) and LYZ2 has been shown to act as an injury-specific marker of pathological extracellular matrix (ECM) degradation (Fan et al., 2025). In addition, gene ontology analysis revealed a marked over-representation of terms related to antioxidant and peroxidase activity, with proteomic GSEA analysis identifying ferroptosis as being markedly enriched in IVH, suggesting that ferroptosis is likely the primary injury axis after neonatal IVH. Cross-omic analysis, identifying the most conserved, robust pathway-level shifts in expression, again highlighted ferroptosis as a key mechanism in neonatal IVH. The cellular components “secondary lysosomes” and “phosphatase complexes” both showed marked, significant enrichment at both gene and protein levels. Iron activation of the lysosome has been implicated in ferroptosis activation and regulation (Cañeque et al., 2025; Chen, 2026), whilst phosphatase complex phosphoglycerate mutase 5 (PGAM5) has been shown to protect against ferroptosis in pancreatitis models by upregulating nuclear factor erythroid 2-related factor 2 (NRF2), widely described as the master regulatory of response to oxidative stress (He et al., 2020; Ma et al., 2026). This implication of ferroptosis in cortical damage was further supported by immunostaining, which identified an IVH-induced depletion of cortical glutathione peroxidase 4 (GPX4), a key buffer against ferroptosis (Li et al., 2022).

To further examine the observed depletion in cortical neurons and neurofilament or synaptic proteins, leading edge analysis was then performed to indicate which concordant gene-protein pairs were core effectors or markers of this process. This suggested that the effects of IVH were primarily marked by expression of piccolo (*Pclo* / PCLO), N-methyl D-aspartate receptor subtype 2B (*Grin2b* / GluN2B), and ELKS/RAB6-interacting/cast family member 1 (*Erc1* / ERC1), all of which have previously been linked to either neuronal stress or neurotransmitter transport at the synapse. While PCLO has been linked to ferroptosis in the context of drug susceptibility in multiple cancers (Tang et al., 2024; Wu et al., 2025), both it and ERC1 are likely represented here due to their functions as scaffold proteins in the presynaptic cytomatrix active zone (Fenster & Garner, 2002; Wang et al., 2002). Similarly, GluN2B is associated with synaptic development and plasticity during neurodevelopment (Akashi et al., 2009). However, while previous studies have indicated a potential selective vulnerability of some GABAergic neuron populations to hypoxia-induced ferroptosis, (Yi et al., 2025), there was no significant difference measured for the GABAergic neuron marker GAD67. This suggests that neuronal depletion after IVH is not specifically targeted to this population of cells, and we hypothesise that the presence of GABAergic neuronal signals here is due to an IVH-induced reduction in GABAergic synaptic function instead of a depletion of GABAergic neurons, though further work must be performed to confirm this.

Transcription factor inference suggested that upregulated DEGs in IVH were primarily driven by increased RAR-related orphan receptor A (RORα) and NRF2 (NFE2L2) activity. Increased NRF2 activity has already been suggested here by the enrichment of multiple ferroptosis and antioxidant defence pathways across both omics’ layers, further supporting their involvement as key processes after IVH. RORα has a key role in cortical maturation (Vitalis et al., 2018) and as a hub gene linking inflammatory and metabolic pathways in hypoxic-ischemic encephalopathy (Song et al., 2025). It has also previously been shown to be elevated in the brain of Parkinson’s disease patients, where it has been suggested to play a neuroprotective effect (Al-Zaid et al., 2023), whilst depletion has been noted in the brains of people living with autism spectrum disorder (V. W. Hu et al., 2015). Inversely, downregulated differentially expressed genes were primarily driven by increased SRY-Box Transcription Factor 7 (SOX7) activity, which is usually associated with glioma suppression (Xiuju et al., 2016; T. Zhao et al., 2016), but has been shown to be a regulator of angiogenesis following hypoxic insult in endothelial cells (Klomp et al., 2020).

Gene ontology analysis of decorin-treated IVH identified over-representation of the “omegasome” gene ontology through increased expression of *Trim12a*, *Trim30d* and *Trim12c,* homologues of TRIM5 which are associated with autophagy and mitophagy (Hatakeyama, 2017; Saha et al., 2022). Decorin has previously been linked to both processes, especially in endothelial cells (Neill et al., 2013), with decorin treatment leading to increases in autophagy (Mishra et al., 2025) and knockdown markedly impairing autophagy *in vivo* (Gubbiotti et al., 2015).

Previous studies have shown that decorin prevents the proliferation of macrophages but promotes their activation and protects them from apoptosis (Comalada et al., 2003). Our study supports this by demonstrating an increase in markers of inflammation and phagocytic activation while showing no significant changes in microglia number or influx of macrophages in the cortex. (Bhide et al., 2005). While decorin can act to reduce inflammation through sequestration of TGF-β1, soluble decorin has also previously been shown to increase inflammatory cytokine production by acting as a ligand to Toll-like receptors 2 and 4 (TLR2 and TL4, respectively) (Merline et al., 2011). Inflammatory response enrichment was well conserved across both omics’ layers even though global transcriptomic-proteomic concordance was lost between IVH and decorin-treated IVH animals. TF inference suggested this was largely regulated by IRF8, IRF4 and STAT1. These TFs are all linked to immune cell regulation, especially of microglia, macrophages, and T cells (Huber & Lohoff, 2014; Moorman et al., 2022; Y. Zhao et al., 2022), with *Irf8* having an important role in determining postnatal microglia identity (Saeki et al., 2024). Therefore, further analysis of the inflammatory response should be performed to fully establish the risk-benefit ratio of using decorin to treat neonatal IVH.

This study is limited by the underrepresentation of some proteins within the LC-MS/MS proteomics, especially high molecular weight ECM proteins and low abundance membrane proteins, as peptide recovery typically biases against these. While our studies included neonates with IVH, neurodevelopmental outcomes may be influenced by factors such as IVH grade, sex, and age. Future research will explore the effects of decorin at other time points, on long-term survival, neurobehavioral outcomes, and sex-based differences. We also examine a single 24-hour post injury snapshot of IVH at PND4, including responses to decorin treatment, which may miss dynamic expression events at both omics’ levels. IVH pathophysiology is partly driven by an immunological response to blood and blood products; therefore, the timing of injury may be critical in influencing severity and treatment efficacy due to differences in immune system maturation at different developmental stages. Indeed, previous work has shown that PND2 IVH rats exhibited minimal activation of the innate immune system compared to those which experienced an IVH at PND5 (Zamorano et al., 2023). Future work may utilise a range of timepoints, ages, and treatment regimens to identify temporal differences in protein and RNA expression.

In conclusion, we describe here the first study of the effects of neonatal IVH on cortical tissues, examining the effects of the proteoglycan decorin when utilised as a treatment for this condition. We demonstrated substantial cortical neurodegeneration following neonatal IVH and identify a marked ferroptosis-associated injury response, both of which are attenuated by decorin treatment. Decorin disrupted the concordant, cross-omics injury signals induced by IVH, supporting the potential of decorin as a potential neuroprotective therapy in this condition, whilst potentially contributing to classical inflammatory signalling. These findings expand current understanding of IVH pathology beyond the ventricular and periventricular compartments, while also supporting the need to consider developmental stage when evaluating injury mechanisms and therapeutic responses. An improved understanding of these processes may enable the development of targeted and specific pharmacological therapies to reduce neuronal death secondary to blood exposure, ultimately working towards reducing neurological disability and improving quality of life in survivors of neonatal IVH.

## 6. Disclosures

### 6.1. Ethics approval and consent to participate

N/A

### 6.2. Availability of data and materials

The RNA-sequencing datasets presented in this study have been deposited at the NCBI sequence read archive (SRA) with BioProject ID PRJNA1415511. The mass spectrometry proteomics data have been deposited to the ProteomeXchange Consortium via the PRIDE (Perez-Riverol et al., 2025) partner repository with the dataset identifier PXD076208. Image analysis macros are available on GitHub (https://github.com/Dr-ben-hewitt/Neonatal-IVH-decorin)

All other data are available upon reasonable request to the corresponding authors.

### 6.4. Authors contributions (CRediT)

BJH: Conceptualisation, Methodology, Software, Formal analysis, Investigation, Visualisation, Writing – Original Draft, Writing – Review and Editing, Funding acquisition, Project administration.

MGB: Methodology, Investigation, Supervision, Writing – Original Draft, Writing – Review and Editing.

JPMcA: Methodology, Supervision, Writing – Review and Editing.

SP: Investigation, Writing – Review and Editing.

GP: Resources, Investigation, Formal analysis, Methodology, Writing - Review and Editing.

MN: Resources, Investigation, Formal analysis, Methodology,

LR: Investigation, Writing - Review and Editing.

JAR: Methodology, Writing - Review and Editing.

PK: Writing - Review and Editing, Supervision.

RMB: Writing - Review and Editing, Supervision.

LJH: Writing - Review and Editing, Supervision.

HFB: Conceptualisation, Methodology, Formal analysis, Investigation, Writing – Original Draft, Writing – Review and Editing, Funding acquisition, Project administration, Supervision.

### 6.6. Acknowledgments

The authors thank the members of the Limbrick Neurosurgery Research Laboratory and Division of Animal Resources (both Virginia Commonwealth University) for their support in this research. We gratefully acknowledge the support of William Bernhardt (Virginia Commonwealth University) with sample shipping. We also thank the Birmingham Environment for Academic Research (BEAR) advanced research computing team for use of high-performance computing services and storage. GP acknowledges Dr Ivana Milic (AIME) for technical assistance.

The Galaxy server used for some calculations is partly funded by the German Federal Ministry of Education and Research BMBF grant 031 A538A de.NBI-RBC and the Ministry of Science, Research and the Arts Baden-Württemberg (MWK) within the framework of LIBIS/de.NBI Freiburg.

