## Supplemental materials for "Cross-omics profiling reveals cortical neuronal dysfunction following neonatal intraventricular haemorrhage that is attenuated by decorin treatment"

**Table S1 – RNA concentration and quality for included samples, as assessed by Nanodrop spectrophotometry.**

| Sample ID | Treatment group | RNA concentration (ng / µl) | RNA quality Abs 260 / 280 nm |
|---|---|---|---|
| A1 | aCSF control | 48 | 2.03 |
| A2 | aCSF control | 1560 | 1.91 |
| A3 | aCSF control | 1149.8 | 2.06 |
| A4 | aCSF control | 943.9 | 2.04 |
| A5 | aCSF control | 909.7 | 2.05 |
| B1 | IVH | 877 | 2.07 |
| B2 | IVH | 781.4 | 2.07 |
| B3 | IVH | 677.1 | 2.05 |
| B4 | IVH | 504.9 | 2.08 |
| B5 | IVH | 793.6 | 2.06 |
| B7 | IVH | 861.9 | 2.06 |
| B9 | IVH | 1108 | 2.07 |
| B10 | IVH | 548.9 | 2.09 |
| B12 | IVH | 632 | 2.08 |
| B13 | IVH | 663.4 | 2.07 |
| C1 | IVH + decorin | 976.7 | 2.08 |
| C2 | IVH + decorin | 733.7 | 2.08 |
| C3 | IVH + decorin | 500.1 | 2.07 |
| C4 | IVH + decorin | 1049.3 | 2.08 |
| C5 | IVH + decorin | 1000.4 | 2.06 |
| C7 | IVH + decorin | 537.3 | 2.09 |
| C9 | IVH + decorin | 616 | 2.08 |
| C10 | IVH + decorin | 690.4 | 2.09 |
| C13 | IVH + decorin | 770.3 | 2.08 |
| C14 | IVH + decorin | 129.9 | 2.04 |

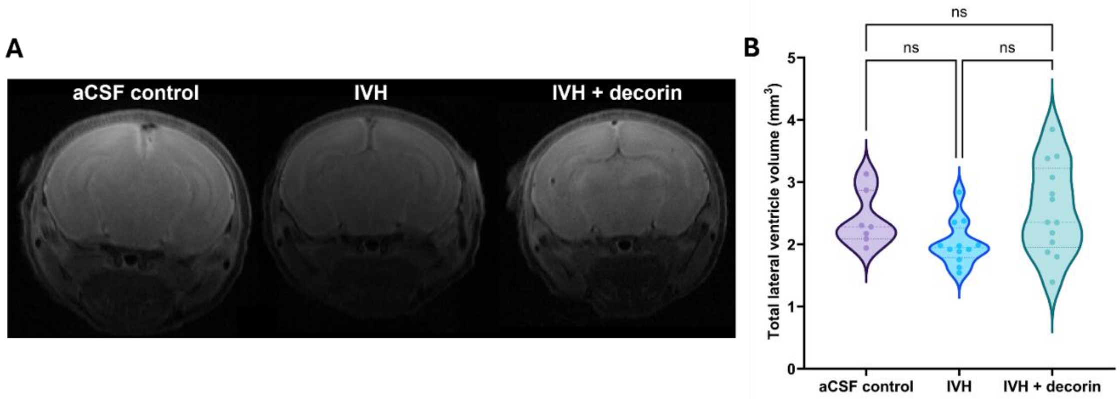


**Figure S1 - Representative MRI images and quantification of ventricular volume.** A) representative MRI sections for aCSF control (N = 7), IVH (N = 12) and IVH + decorin (N = 13) groups. B) Violin plot of quantified lateral ventricle volume (left and right combined) for each animal. Statistical testing performed with one-way ANOVA followed by Tukey's multiple comparisons test, ns = not significant, α = 0.05.

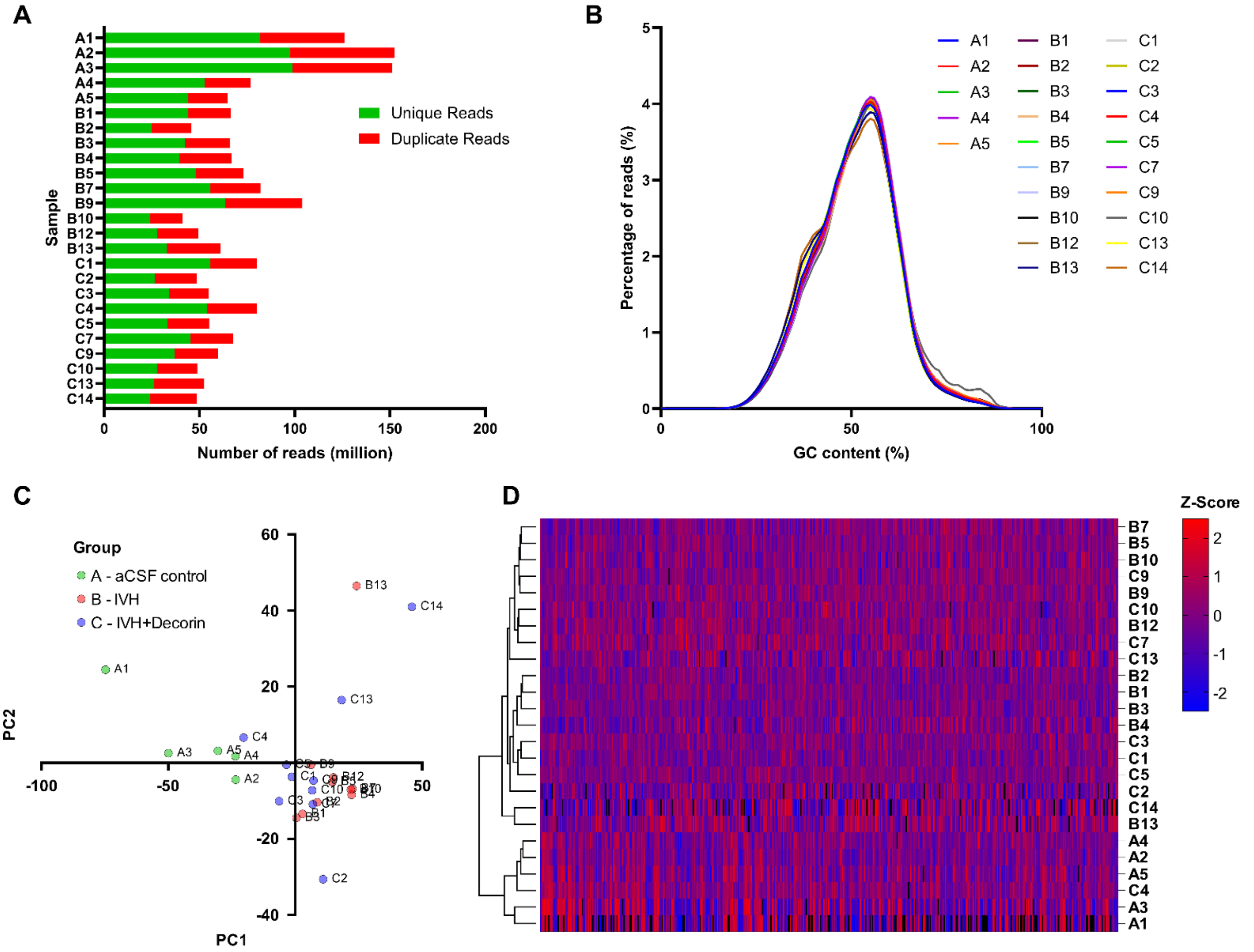


**Figure S2 - Quality control and clustering analyses for aCSF control (N = 5), IVH (N = 10) and IVH+decorin (N = 10) groups (groups A, B and C respectively).** A) Read mapping quality control showed an acceptable number of duplicate reads, with each sample having a minimum mapping of 20 million unique reads / sample. B) GC content graph as a function of number of reads, showing a mean GC content of ~50%. C) PCA of variance stabilised expression (VST) scores from top 500 differentially expressed genes (DEGs), showing distinct clustering of aCSF control samples from IVH and IVH + decorin, but overlap of IVH and IVH + decorin groups. D) Hierarchical clustering on top 500 DEGs, showing distinct clustering of aCSF control and IVH / IVH + decorin animals. Euclidean distance method, columns standardised and represented as Z-score (red = upregulation, blue = downregulation).

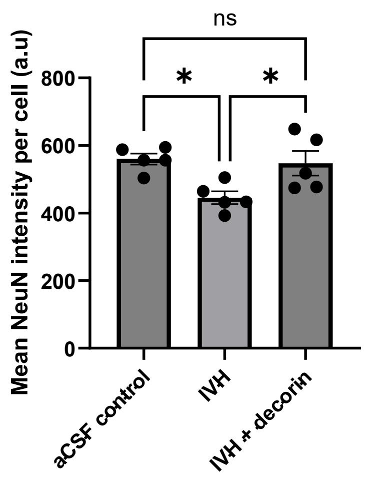


**Figure S3 – Mean NeuN intensity of NeuN staining per cell.** One way ANOVA with Tukey's multiple comparisons test (all to all). N = 5 biological replicates (no outliers). * = *P* < 0.05, ns = *P* > 0.05.

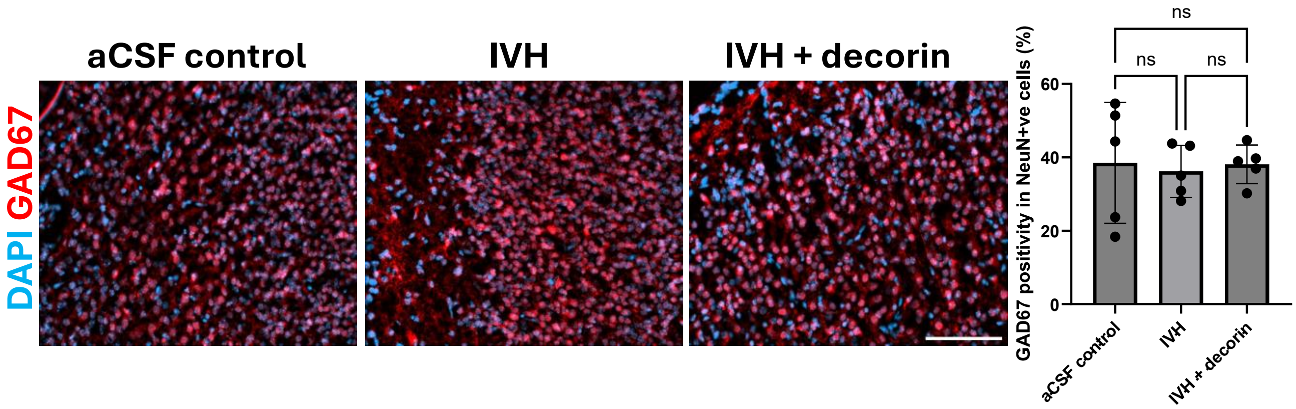


**Figure S4 – GAD67 immunostaining in the cortex.** One way ANOVA with Tukey's multiple comparisons test (all to all). ns = $P > 0.05$.

**Table S2 – Proteins utilised in hypothesis-driven analysis of the effects of IVH and decorin treatment on ferroptosis, iron handling, redox regulation, neurotransmitter release and neuronal structural or axonal proteins.**

| Protein | Panel | Rationale | Reference |
|---|---|---|---|
| FTH1, FTL1, TF, TFRC | Ferroptosis and iron handling | Iron sequestration | (Dixon & Stockwell, 2018) |
| FADS2, ACOX1 | Ferroptosis and iron handling | Lipid metabolism | (Hu et al., 2025; Yamane et al., 2022) |
| PRDX6, PRDX1, PRDX2, GSTP1, GSTM1, PARK7, SOD2 | Antioxidant related | Antioxidants and redox regulation | (Shim & Kim, 2013; Tew & Townsend, 2012) |
| SNAP25, STX1A, STX1B, SYT1, VAMP2 | Synapse-related | Neurotransmitter release | (Südhof, 2013) |
| DNM1, RAB3a | Synapse-related | Vesicle trafficking | (Ferguson & De Camilli, 2012; Südhof, 2004) |
| GAP43 | Neuronal structural and axonal proteins | Axon growth | (Benowitz & Routtenberg, 1997) |
| MAP2 | Neuronal structural and axonal proteins | Neurogenesis | (Dehmelt & Halpain, 2005) |
| NEFL, NEFM, INA | Neuronal structural and axonal proteins | Neurofilament | (Yuan et al., 2017) |
| TUBB3 | Neuronal structural and axonal proteins | Axon guidance | (Puri et al., 2023) |
